# A bacterial derived plant- mimicking cytokinin hormone regulates social behaviour in a rice pathogen

**DOI:** 10.1101/2021.07.05.451090

**Authors:** Sohini Deb, Chandan Kumar, Rahul Kumar, Amandeep Kaur, Palash Ghosh, Gopaljee Jha, Prabhu B. Patil, Subhadeep Chatterjee, Hitendra K. Patel, Ramesh V. Sonti

**Affiliations:** CSIR- Centre for Cellular and Molecular Biology (CSIR-CCMB), Hyderabad- 500007, India; National Institute of Plant Genome Research (NIPGR), New Delhi- 110067, India; CSIR - Institute of Microbial Technology (CSIR-IMTECH), Chandigarh- 160036, India; Centre for DNA Fingerprinting and Diagnostics (CDFD), Hyderabad- 500039, India; Indian Institute of Science Education and Research, Tirupati-517507, India

**Keywords:** *Xanthomonas*, type III effector, biofilm, cytokinin

## Abstract

Many plant-associated bacteria produce plant- mimicking hormones which are involved in modulating host physiology. However, their function in modulating bacterial physiology has not been reported. Here we show that the XopQ protein, a type-III effector of the rice pathogen, *Xanthomonas oryzae* pv. *oryzae* (*Xoo*), is involved in cytokinin biosynthesis. *Xoo* produces and secretes an active form of cytokinin which enables the bacterium to maintain a planktonic lifestyle and promotes virulence. RNA-seq analysis indicates that the cytokinin produced by *Xoo* is required for the regulation of several genes which are involved in biofilm formation. We have also identified the *Xoo* isopentenyl transferase gene, which is involved in the cytokinin biosynthesis pathway and is required for maintaining planktonic behaviour and virulence. Furthermore, mutations in the predicted cytokinin receptor kinase (PcrK) and the downstream response regulator (PcrR) of *Xoo* phenocopy the cytokinin biosynthetic mutants, but are not complemented by supplementation with exogenous cytokinin. Cytokinin biosynthetic functions are encoded in a number of diverse bacterial genomes suggesting that cytokinin may be a widespread signalling molecule in the bacterial kingdom.

## Introduction

Cytokinins are plant hormones that promote various aspects of plant growth, development and immunity (Osugi & Sakakibara, 2015). Several plant pathogenic bacteria such as *Rhodococcus fascians, Agrobacterium tumefaciens* and *A. rhizogenes* strains have been shown to produce cytokinins as part of their virulence repertoire in order to modulate host physiology (Pertry, Václavíková et al., 2009, Sardesai, Lee et al., 2013). Cytokinin production by the plant growth promoting bacterium *Pseudomonas fluorescens* and the presence of intact plant cytokinin receptors has been shown to be necessary for biocontrol activity in *Arabidopsis thaliana* (Großkinsky, Tafner et al., 2016). *Mycobacterium tuberculosis* has also been shown to encode a phosphoribose-hydrolase that converts isopentenyl adenosine monophosphate (iPMP) to isopentenyl adenine (iP) and that it accumulates iP and 2-methylthio-iP in the culture medium (Samanovic, Tu et al., 2015). However, it is not known why *M. tuberculosis* produces cytokinin. *Corynebacterium glutamicum* encodes two proteins that can function as phosphoribose-hydrolases and a large number of prokaryotic organisms have been shown to have homologs of these enzymes (Samanovic et al., 2015, Seo & Kim, 2017). This suggests the intriguing possibility that cytokinins may be made by a number of bacteria and that these compounds may have a role in regulating bacterial physiology. Although it is well known that certain bacteria produce cytokinin to regulate host physiology, there is no evidence to date that endogenously produced cytokinin is used by bacteria to modulate their own physiology or cellular behaviour. Recently, a receptor for host produced cytokinin, named as Plant cytokinin receptor Kinase (PcrK), has been identified in the bacterium *Xanthomonas campestris* pv. *campestris* (*Xcc*), which can sense exogenously produced cytokinin. PcrK is a histidine kinase that is a part of the PcrK/PcrR two-component system, activation of which has been shown to enhance bacterial resistance to reactive oxygen species, produced as a part of the host defense response (Wang, Cheng et al., 2017).

The *Xanthomonas oryzae* pv. *oryzae* (*Xoo*) type III effector Xanthomonas outer protein Q (XopQ) is a homolog of the *Pseudomonas syringae* type III effector HopQ1, which appears to have phosphoribose-hydrolase activity (Hann, Dominguez-Ferreras et al., 2014). Here we report that XopQ is a phosphoribose-hydrolase which acts on the cytokinin precursor iPMP to produce cytokinin. Endogenous cytokinin production appears to control the ability of the bacterium to remain in a planktonic state as the *xopQ-* mutant shows a tendency to form aggregates and enhanced biofilm formation. External supplementation of cytokinin to the *xopQ-* mutant restores its ability to remain in a planktonic mode as well as complements its virulence deficiency. We have further identified the *Xoo* isopentenyl transferase (*ipt*) gene which catalyzes an earlier step in the cytokinin biosynthetic pathway in *Xoo*. Mutation in the *ipt* gene predisposes the bacterium to form aggregates, enhances biofilm formation and reduces virulence; all of which can be restored by external supplementation of cytokinin. Thus, the *ipt-* mutant mimics the *xopQ-* mutant, consistent with the observation that they both affect cytokinin biosynthesis. RNA-seq analysis indicates differential expression of a number of genes in the *xopQ-* mutant that can affect biofilm formation.

## Results

### The XopQ protein is a phosphoribose-hydrolase which produces cytokinin enabling the planktonic growth of *X. oryzae* pv. *oryzae*

Earlier work had indicated that the aspartate residue at 116^th^ position and tyrosine at 279^th^ position are important for the phosphoribose-hydrolase activity of XopQ (Gupta, Nathawat et al., 2015). Hence, the purified recombinant proteins XopQ, XopQ D116A and XopQ Y279A were assayed for activity using the putative substrate isopentenyl adenosine monophosphate (iPMP). Activity of the wildtype XopQ protein, as well as mutant proteins, was found to increase with increasing substrate concentration but did not reach saturation. Activity was found to decrease at higher concentrations (Appendix Figure S1); possibly due to substrate inhibition. Hence, kinetic parameters were calculated using the substrate concentrations at which a linear increase in activity was observed. Similar K_m_ values for the wildtype and mutant indicated that mutations in the putative catalytic site of XopQ do not affect the affinity of the XopQ enzyme for the substrate (Table 1, Appendix Figure S1). However, the K_cat_ values indicated a strong activity of XopQ toward iPMP, as well as a significant reduction in the activity of the mutant proteins as compared to the wildtype XopQ protein. The lower K_cat_/K_m_ value for the mutant proteins as compared to the wildtype XopQ protein suggested that the mutant proteins have less catalytic efficiency than the wildtype protein towards iPMP.

**Table 1.** The XopQ protein is a phosphoribose-hydrolase which cleaves a cytokinin precursor. Kinetic parameters of phosphoribose-hydrolase activity of XopQ wildtype and mutant proteins XopQ D116A and XopQ Y279A of *Xoo*.

| Parameters | XopQ | XopQ D116A | XopQ Y279A |
| --- | --- | --- | --- |
| $V_{\max}$ ($\mu\text{M sec}^{-1}$) | 2.84808E-05 | 2.2531E-05 | 1.96002E-05 |
| $K_m$ ($\mu\text{M}$) | 0.191494727 | 0.174519165 | 0.180779251 |
| $K_{\text{cat}}$ ($\text{sec}^{-1}$) | 28.4808237 | 23.0377589 | 19.6002059 |
| $K_{\text{cat}}/K_m$ ($\text{M}^{-1} \text{sec}^{-1}$) | 0.000148729 | 0.000132007 | 0.000108421 |

XopQ transcripts and protein were found to be expressed in the wildtype *Xoo* strain BXO43 in laboratory PS media (Appendix Figure S2A, B). In order to determine if BXO43 produces cytokinin in laboratory cultures, we used LC/MS to estimate the amounts of the two cytokinins, isopentenyl adenine (iP) and trans-zeatin (tZ), in the cell pellet and culture supernatant of BXO43, mutant (*xopQ-*) and complement strains (*xopQ-*/pHM1, *xopQ-* /pHM1::*xopQ, xopQ-*/pHM1::*xopQ D116A* and *xopQ-*/pHM1::*xopQ Y279A*). Interestingly, the amount of iP produced by BXO43 in both cell pellet and supernatant was nearly 100-fold higher as compared to tZ (Fig 1A-D). As compared to BXO43, the *xopQ-* mutant produced significantly lesser amount of both iP as well as tZ, which could be complemented by introduction of the *xopQ* wildtype gene into the *xopQ-* strain through the pHM1 vector. Notably, the *xopQ D116A* and *xopQ Y279A* mutants failed to complement the reduction in cytokinin production of the *xopQ-* strain (Fig 1A-D).

**Figure 1.**
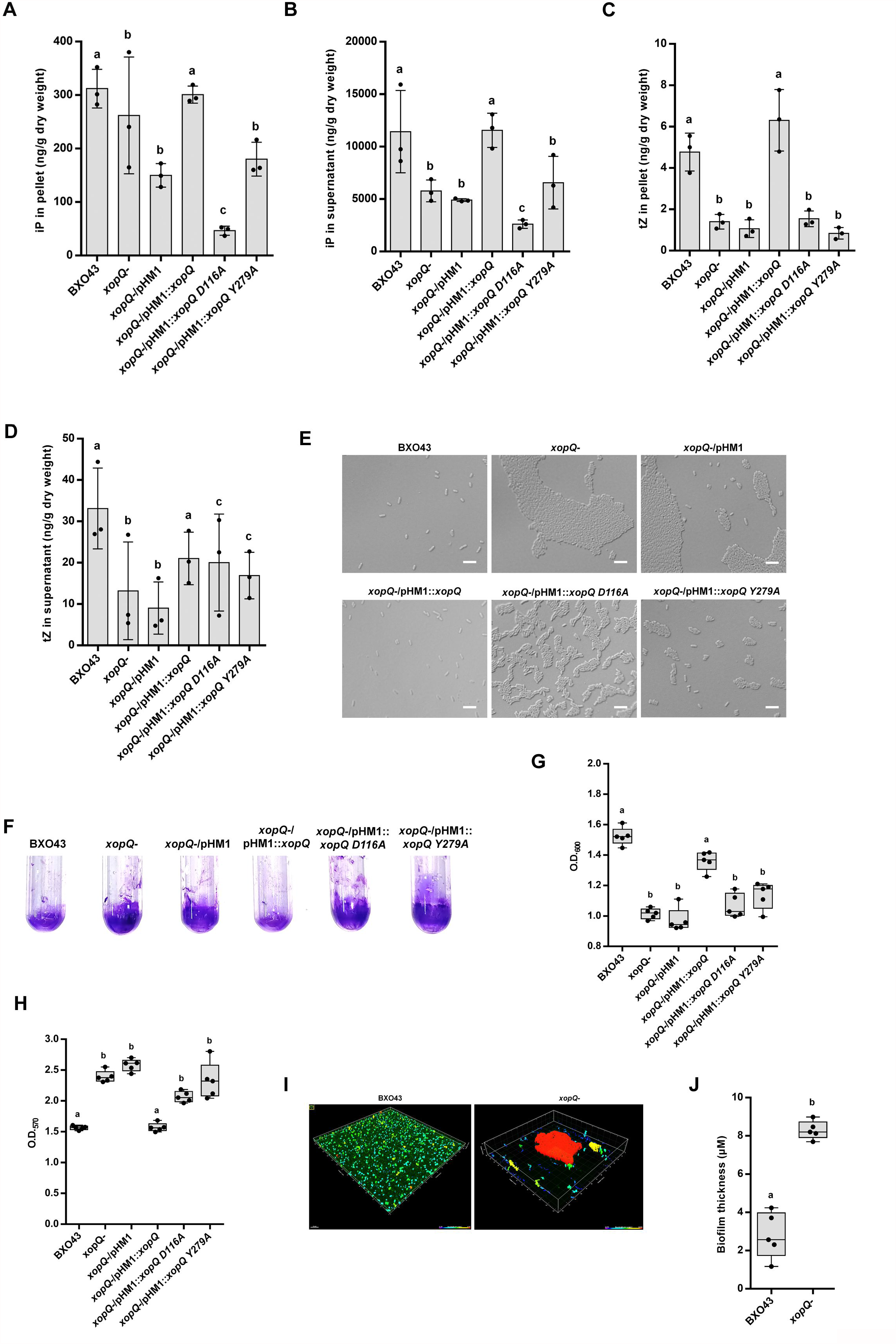
The XopQ protein is a phosphoribose-hydrolase which produces and secretes cytokinin for the planktonic growth of *X. oryzae* pv. *oryzae*. **A-D**, Cytokinin estimation was carried out from the bacterial cell pellets and culture supernatant by LC-MS using the different *Xoo* strains. Isopentenyl adenine (iP) (**A, B**) and trans-zeatin (tZ) (**C, D**) was measured in cell pellet and culture supernatant. Values are presented as nanogram of cytokinin per gram of dry weight of lyophilised sample ± standard deviation from 3 biological replicates. **E**, Microscopy images of the different *Xoo* strains. Images were acquired at 100X in DIC (Nomarski). Scale bar represents 5 µm. **F-H**. Cell attachment and static biofilm assay of *Xoo* strains. **G**, Quantification of bacterial cells in the cell suspension. Values are presented as mean absorbance (at 600 nm) ± standard deviation from 5 replicates. **H**, Quantification of bacterial cells attached to glass tubes by staining with crystal violet. Values are presented as mean absorbance (at 570 nm) ± standard deviation from 5 replicates. **I-J**, Surface visualisation of biofilm formed by the BXO43/EGFP and *xopQ-*/EGFP strains. Scale bar represents 15 µm. Thickness of the biofilm is presented as mean ± standard deviation from 5 replicates. Asterisk indicates significant difference (*P*= 0.0197) in comparison with the BXO43 strain. (**J**). For all graphs, columns/boxes capped with letters that are different from one another indicate that they are statistically different using unpaired two-sided Student’s t-test analysis (*P* ≤ 0.05). Images are representative of 3 biological replicates (**E, F, I**).

Microscopic analysis revealed that *xopQ-* cells tend to form aggregates, in comparison to BXO43 cells, which remain dispersed (Fig 1E). We further went on to visualise the complement strains of *xopQ-*, i.e., *xopQ-*/pHM1, *xopQ-*/pHM1::*xopQ, xopQ-*/pHM1::*xopQ D116A* and *xopQ-*/pHM1::*xopQ Y279A*. The *xopQ-*/pHM1 strain formed aggregates, similar to the *xopQ-* mutant. However, complementation of the *xopQ-* cells with the wildtype *xopQ* gene restored a planktonic mode. Mutants in the *xopQ* gene which affected biochemical activity and cytokinin production, namely the *xopQ-*/pHM1::*xopQ D116A* or *xopQ-* /pHM1::*xopQ Y279A* strains, formed aggregates similar to the *xopQ-* strain (Fig 1E).

We reasoned that the ability to form aggregates may result in a higher ability to form biofilms. To examine the role of *xopQ* in attachment and biofilm formation, we performed quantitative cell attachment and static biofilm assays using glass test tubes. The BXO43 strain was seen to exhibit a minimal amount of biofilm formation as assayed after 4 days under biofilm formation conditions. However, the *xopQ-* strain formed significantly more biofilm, as visualized by staining with crystal violet (Fig 1F-H). This was also visualised by using the BXO43 or *xopQ-* strains expressing EGFP on a plasmid (Fig 1I, J). Introduction of the pHM1 empty vector into the *xopQ-* mutant did not alter the biofilm formation phenotype of the *xopQ-* strain. However, introduction of the wildtype *xopQ* gene on the complementing plasmid resulted in reduction in biofilm formation, a phenotype that was similar to that of BXO43. The biochemically inactive mutants *xopQ D116A* or *xopQ Y279A* were like the *xopQ-* mutant (Fig 1F-H).

### Supplementation with exogenous cytokinin converts a *xopQ-* mutant from biofilm to planktonic phenotype and restores wildtype levels of virulence

We examined whether addition of exogenous cytokinin would rescue the aggregation phenotype of the *xopQ-* cells. For this, the cytokinin iP was added to actively growing cultures of either BXO43 or the *xopQ-* mutant. Addition of iP could disperse the aggregates formed by cells of *xopQ-* (Fig 2A). Surprisingly, addition of cytokinin induced aggregate formation in the BXO43 strain. We also examined the ability of these strains to form biofilm. As described previously, the *xopQ-, xopQ-*/pHM1, *xopQ-*/pHM1::*xopQ D116A* and *xopQ-* /pHM1::*xopQ Y279A* strains formed more biofilm as compared to BXO43 or the *xopQ-* /pHM1::*xopQ* strains (Fig 1F-H). When iP was added, biofilm formation by the *xopQ-, xopQ-* /pHM1, *xopQ-*/pHM1::*xopQ D116A* and *xopQ-*/pHM1::*xopQ Y279A* strains reduced significantly (Fig 2B-D). Addition of cytokinin induced higher biofilm formation in the BXO43 and *xopQ-*/pHM1::*xopQ* strains. Addition of iP to cultures of EGFP expressing derivatives of BXO43 and *xopQ-* strains led to reduced biofilm formation by the *xopQ-* strain, whereas it enhanced biofilm formation by BXO43 (Fig 2E-F).

**Figure 2.**
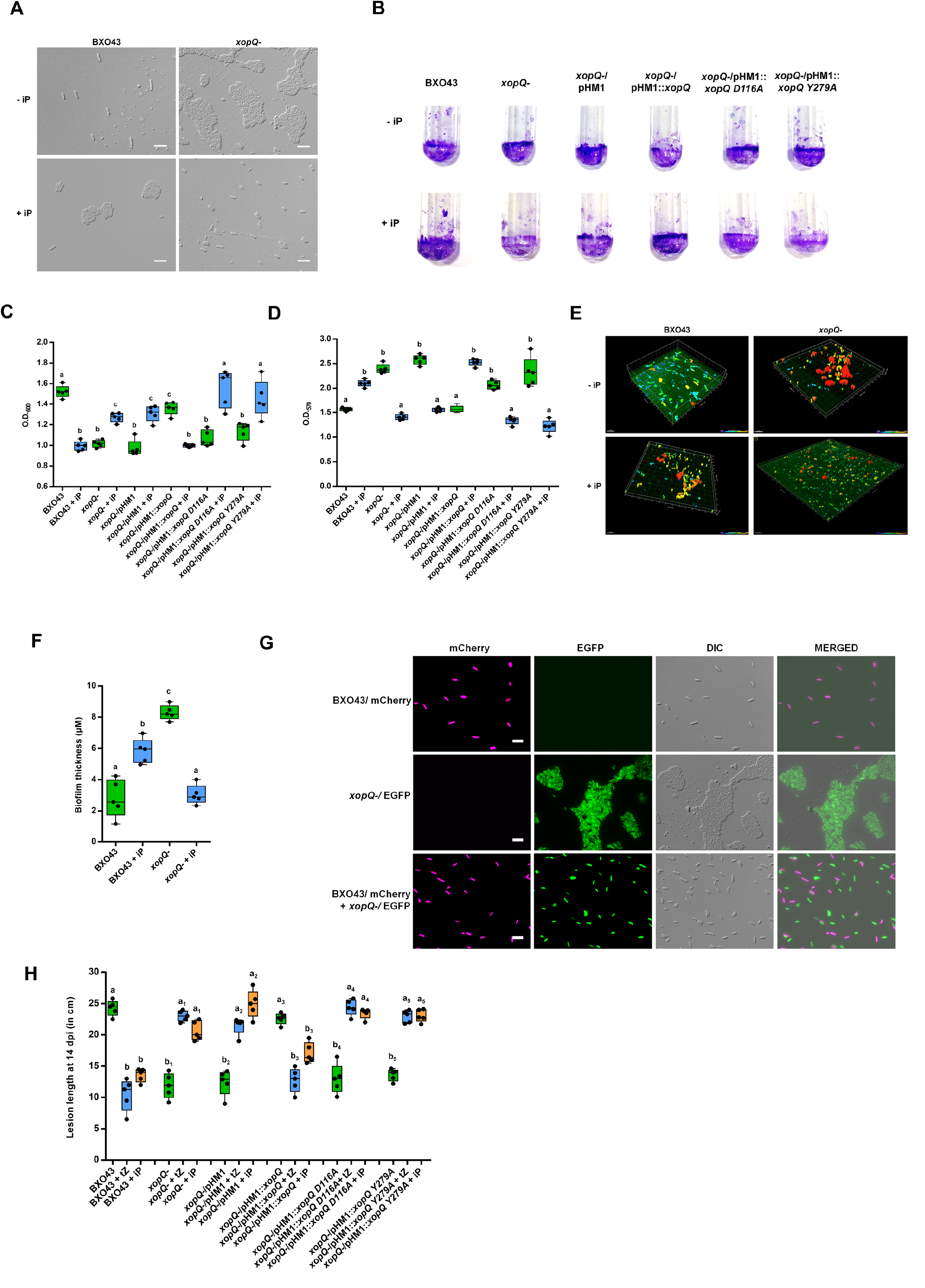
Supplementation with exogenous cytokinin converts a *xopQ-* mutant from biofilm to planktonic lifestyle and restores wildtype levels of virulence. **A**, Microscopy analysis was performed of the *Xoo* strains, with and without the addition of iP. Images were acquired at 100X in DIC (Nomarski). Scale bar represents 5 µm. **B-D**. Cell attachment and static biofilm assay was performed of the *Xoo* strains, with and without the addition of iP. **C**, Quantification of bacterial cells in the cell suspension. Values are presented as mean absorbance (at 600 nm) ± standard deviation from 5 replicates. **D**, Quantification of bacterial cells attached to glass tubes by staining with crystal violet. Values are presented as mean absorbance (at 570 nm) ± standard deviation from 5 replicates. **E-F**, Surface visualisation of biofilm formed by the BXO43/EGFP and *xopQ-*/EGFP strains, with and without the addition of iP. Scale bar represents 15µm. **F**, Thickness of the biofilm is presented as mean ± standard deviation from 5 replicates. **G**, Co- culturing of wildtype *Xoo* BXO43 with the *xopQ-* mutant reverses biofilm to planktonic lifestyle of *Xoo*. Microscopy analysis was performed of the BXO43/mCherry and *xopQ-*/EGFP strains, either singly, or following co- culturing for 4 h at 28 °C. Images were acquired at 100X for fluorescence channels and DIC (Nomarski). Scale bar represents 5 µm. **H**, Supplementation with exogenous cytokinin restores wildtype levels of virulence to a *xopQ-* mutant of *Xoo*. Rice leaves were inoculated by pin- pricking with the various *Xoo* strains, with and without prior midvein injection of iP. Lesion lengths were measured 14 days after inoculation. Error bars indicate the standard deviation of readings from 5 inoculated leaves. For all graphs, boxes capped with letters that are different from one another indicate that they are statistically different using unpaired two- sided Student’s t-test analysis (*P* ≤ 0.05). Images are representative of 3 biological replicates (**A, B, E, G**).

We also examined if co-culture with the BXO43 strain would rescue the aggregate formation phenotype of the *xopQ-* strain. In order to distinguish the BXO43 and *xopQ-* strains, a *P*_*lac*_*_*mCherry plasmid was introduced into BXO43, while a EGFP plasmid was introduced into the *xopQ-* strain. When cultured individually, BXO43/mCherry cells were dispersed, whereas *xopQ-*/EGFP cells formed aggregates. On co-culturing these two strains, we observed that the *xopQ-*/EGFP strain no longer formed aggregates, and appeared to be dispersed (Fig 2G). These results suggest that, during co-culture, the cytokinin secreted by the BXO43 strain can rescue the aggregation phenotype of the *xopQ-* mutant.

The *xopQ-, xopQ D116A* and *xopQ Y279A* mutants of *Xoo* exhibit a virulence deficiency *in-planta* (Gupta et al., 2015). We determined whether external supplementation with active forms of cytokinin such as tZ or iP would restore the virulence deficiency of the *xopQ-* mutant. For this purpose, the BXO43, *xopQ-, xopQ-*/pHM1, *xopQ-*/pHM1::*xopQ, xopQ-* /pHM1::*xopQ D116A* and *xopQ-*/pHM1::*xopQ Y279A* strains were assayed for virulence on rice with or without addition of tZ or iP. In the absence of cytokinin, lesion lengths formed by either BXO43 or wildtype *xopQ* complemented strain were significantly longer than those obtained after infection with *xopQ-* or *xopQ-* expressing pHM1 vector alone, or pHM1 expressing *xopQ D116A* or *xopQ Y279A* (Fig 2H). However, in the presence of tZ or iP, virulence of *xopQ-, xopQ-*/pHM1::*xopQ D116A* or *xopQ-*/pHM1::*xopQ Y279A* strains was restored to wildtype levels. These observations indicate that supplementation with exogenous cytokinin restores wildtype levels of virulence to the *xopQ-* mutant. Surprisingly, addition of cytokinin in the presence of the wildtype copy of *xopQ* (i.e., BXO43 or *xopQ-*/pHM1::*xopQ*), resulted in reduced virulence of these strains, suggesting that an optimum level of cytokinin is necessary for complete virulence of *Xoo* (Fig 2H).

### The isopentenyl transferase gene is required for planktonic lifestyle and full virulence of *Xoo*

The isopentenyl transferase (*ipt*) gene encodes the committed step in the biosynthetic pathway of cytokinins. Putative IPT proteins were identified in *X. theicola, X. axonopodis, X. bromi, X. albilineans, X. translucens, X. oryzae* pv. *oryzae* PXO99a, *X. oryzae* pv. *oryzae* BXO1 and *X. oryzae* pv. *oryzicola* BLS256 by using *Agrobacterium tumefaciens* IPT protein as a query in NCBI GenBank. Alignment of the protein sequences revealed a high degree of conservation of this protein among these *Xanthomonas* species (Appendix Figure S3A). However, the *ipt* gene was absent in a few *Xanthomonas* species such as *X. campestris* pv. *vesicatoria* (*Xcv*) and *X. campestris* pv. *campestris* (*Xcc*) which infect dicotyledonous plants. Further bioinformatics analysis indicated that along with *ipt*, a 4777 bp region encompassing *ipt* is absent in both *Xcv* and *Xcc* (Fig 3A).

**Figure 3.**
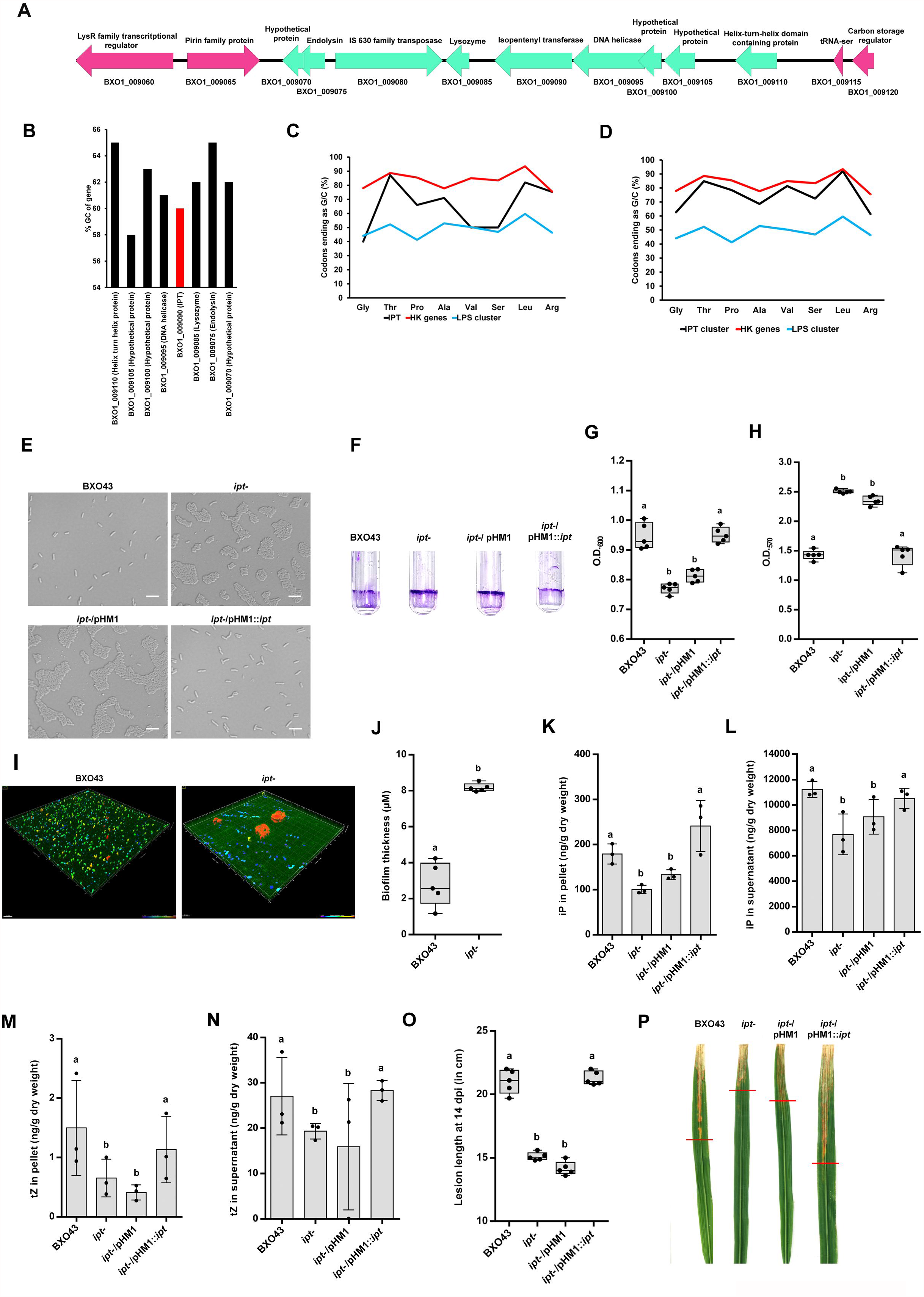
The isopentenyl transferase gene is required for planktonic lifestyle and full virulence of *Xoo*. **A**, Schematic of open reading frames (ORF) based on sequence of 4777 bp genomic region encompassing the IPT locus of the *Xoo* BXO1 strain. Arrows represent the ORF and direction of transcription. The predicted ORFs upstream of *ipt* gene encode a hypothetical protein, an endolysin, an IS630 transposase and a lysozyme. The predicted ORFs downstream of *ipt* gene in BXO1 exhibit high similarity to a DNA helicase and a helix-turn-helix protein. ORFs marked in green represent the 4777 bp region present in *Xoo* but not in *Xcv* or *Xcc*. ORFs marked in pink denote the flanking genes, conserved in *Xcv* and *Xcc* **B**, GC content of IPT locus. **C**, Codon usage pattern of *ipt* gene, HK (housekeeping) genes and LPS cluster **D**, Codon usage pattern of *ipt* gene cluster (excluding IS630 transposase), HK genes and LPS cluster **e**, Microscopy analysis was performed of the *Xoo* strains. Images were acquired at 100X in DIC (Nomarski). Scale bar represents 5 µm. **F-H**, Cell attachment and static biofilm assay of *Xoo* strains BXO43, *ipt-, ipt-*/pHM1 or *ipt-*/pHM1::*ipt*. **G**, Quantification of bacterial cells in the cell suspension. Values are presented as mean absorbance (at 600 nm) ± standard deviation from 5 replicates. **H**, Quantification of bacterial cells attached to glass tubes by staining with crystal violet. Values are presented as mean absorbance (at 570 nm) ± standard deviation from 5 replicates. **I-J**, Surface visualisation of biofilm formed by the BXO43/EGFP and *ipt-*/EGFP strains. Scale bar represents 15 µm. **J**, Thickness of the biofilm is presented as mean ± standard deviation from 5 replicates. Asterisk indicates significant difference (*P*= 0.0209) in comparison with the BXO43 strain. **K-N**, Cytokinin estimation was carried out from the bacterial cell pellets and culture supernatant of *Xoo* strains by LC-MS. iP (**K, L**) and tZ (**M, N**) was measured in cell pellet and culture supernatant. Values are presented as nanogram of cytokinin per gram of dry weight of lyophilised sample ± standard deviation from 3 biological replicates. **O-P**, A *ipt-* mutant of BXO43 is virulence deficient. Leaves of susceptible rice TN-1 were clip inoculated with different *Xoo* strains. **O**, Lesion lengths were measured 14 days after inoculation. Error bars indicate the standard deviation of readings from 5 inoculated leaves. **P**, Virulence phenotype on rice leaves. Leaves were photographed 14 days after inoculation. For all graphs, columns/boxes capped with letters that are different from one another indicate that they are statistically different using unpaired two- sided Student’s t-test analysis (*P* ≤ 0.05). Images are representative of 3 biological replicates (**E, F, I, P**).

The *ipt* transcripts were found to be expressed in PS medium grown cultures of the BXO43 strain (Appendix Figure S4). Analysis of the *ipt* gene in the *Xoo* genome indicated that it is conserved in nearly 100 sequenced Indian isolates of *Xoo* with 100% coverage and identity (unpublished observations, Prabhu B. Patil). Interestingly, the *ipt* gene has a G+C content of 60%, which is significantly lesser than the average G+C content of *Xoo*, which is 64-65%, suggesting that *ipt* might have been acquired by horizontal gene transfer (Fig 3B). We also examined the codon usage pattern (CUP) of the *ipt* gene and observed that CUP of *ipt* as well of the 4777 bp region was significantly different from that of the housekeeping genes of *Xoo* and of the O-antigen biosynthetic gene cluster of *Xoo*, which has been earlier shown to have the signature features of a genomic island (Patil & Sonti, 2004) (Fig 3C, D). Also, the presence of a IS630 family transposase and a tRNA-ser is consistent with this locus being a genomic island that has been acquired through horizontal gene transfer. This tRNA gene is present at the orthologous location in *Xcc* and *Xcv*, although the entire *ipt* genomic island is lacking in *Xcv* and *Xcc*. Bioinformatics analysis indicates that genes which encode homologs of the IPT protein are encoded in a number of bacteria (Appendix Figure S3B). Phylogenetic analyses of IPT proteins from diverse organisms has revealed the conservation of IPT proteins and their grouping into three clades (Appendix Figure S3B). The first clade contained only bacterial species with two subgroups of plant associated bacteria and soil bacteria; the second clade contained tRNA-type IPTs from all four groups, namely, Archaea, bacteria, fungi and plants, whereas the third group contained adenylate-type IPTs from fungi and plants.

Microscopic analysis of the *ipt-* strain revealed that these cells form aggregates, whereas cells of the wildtype (BXO43) remained dispersed (Fig 3E). Further, complementation of the *ipt-* mutant with the wildtype *ipt* gene reduced aggregate formation and restored the ability of the bacteria to grow in a planktonic mode. The *ipt-* strain also formed more biofilm as compared to BXO43 and complementation with the wildtype *ipt* gene restored the wildtype phenotype (Fig 3F-H). Microscopic analysis of EGFP expressing strains revealed that the *ipt-* strain formed a thicker biofilm as compared to BXO43 (Fig 3I-J). We further estimated the cytokinin content (tZ and iP) in both cell pellet as well as supernatant of the *ipt-* strain. As compared to BXO43, the *ipt-* strain showed a significant reduction in levels of both iP as well as tZ, which could be complemented by introduction of the wildtype *ipt* gene into the *ipt-* strain through the pHM1 vector (Fig 3K-N). The reduction in cytokinin levels, especially for iP, appears to be less than the reduction seen in the *xopQ-* mutant. This suggests that there may be other substrates, besides those produced through action of IPT, on which XopQ can act to produce iP.

We then proceeded to examine the virulence of the *ipt-* strain. Leaves of 60-day old rice plants were inoculated with cultures of BXO43, *ipt-, ipt-*/pHM1 or *ipt-*/pHM1::*ipt*. The *ipt-* strain, as well as *ipt-* carrying the empty vector pHM1 showed a significant reduction in lesion length as compared to BXO43 at 14 days post inoculation. Introduction of the *ipt* gene in the *ipt-* mutant restored wildtype levels of virulence (Fig 3O-P).

### Supplementation of *ipt-* mutant with active cytokinin restores the wildtype phenotype

In order to test if exogenous cytokinin addition would rescue the aggregate formation of an *ipt-* mutant, we added iP to actively growing cultures of the *ipt-* mutant. Addition of iP could disperse the aggregates formed by the *ipt-* mutant (Fig 4A). Cytokinin supplementation to the *ipt-* strain also led to reduced biofilm formation (Fig 4B-F). This was reflected in a lesser density of cells in culture in the *ipt-* strain as compared to BXO43 or *ipt-* + iP (Fig 4C). In order to check if the cytokinin secreted by BXO43 would rescue aggregate formation by *ipt-*, we went ahead to co-culture the BXO43/mCherry strain with the *ipt-*/EGFP strain. When cultured individually, BXO43/mCherry cells were dispersed, whereas *ipt-*/EGFP cells formed aggregates. On co-culturing these two strains, we observed that the *ipt-*/EGFP strain is dispersed and no longer formed aggregates (Fig 4G). This suggests that the cytokinin secreted by the BXO43 strain rescues the aggregation phenotype of the *ipt-* strain.

**Figure 4.**
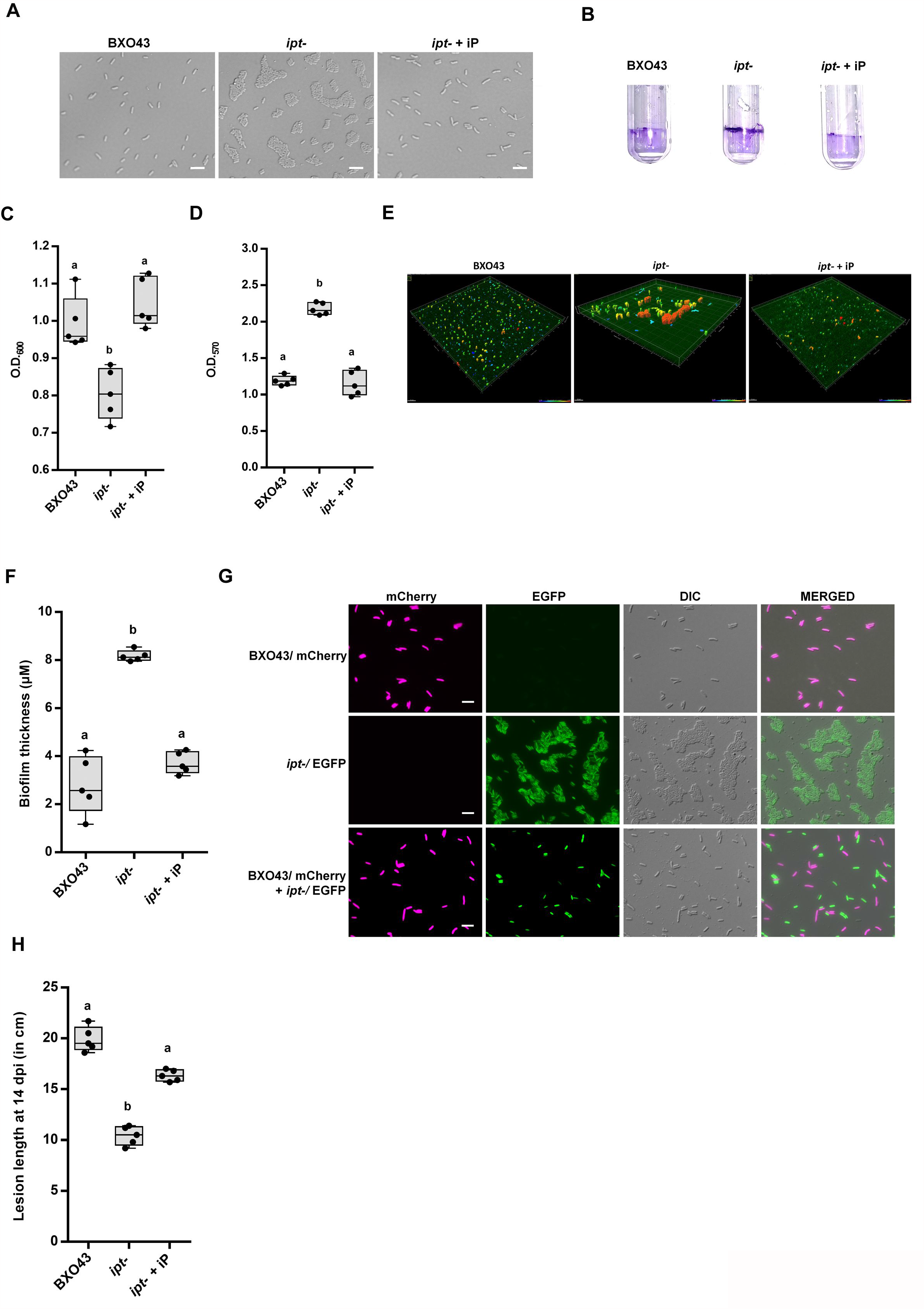
Supplementation of *ipt-* with active cytokinin reverses biofilm to planktonic lifestyle and restores wildtype levels of virulence. **A**, Microscopy analysis was performed of the following strains: BXO43, *ipt-* or *ipt-* + iP. Images were acquired at 100X in DIC (Nomarski). Scale bar represents 5µm. **B-D**. Cell attachment and static biofilm assay of *Xoo* strains BXO43, *ipt-* or *ipt-* + iP. **c**, Quantification of bacterial cells in the cell suspension. Values are presented as mean absorbance (at 600 nm) ± standard deviation from 5 replicates. **D**, Quantification of bacterial cells attached to glass tubes by staining with crystal violet. Values are presented as mean absorbance (at 570 nm) ± standard deviation from 5 replicates. **E- F**, Surface visualisation of biofilm formed by the BXO43/EGFP and *ipt-*/EGFP strains. Scale bar represents 15 µm. **F**, Thickness of the biofilm is presented as mean ± standard deviation from 5 replicates. **G**, Co-culturing of wildtype *Xoo* BXO43 with the *ipt-* mutant reverses biofilm to planktonic lifestyle of *Xoo*. Microscopy analysis was performed of the BXO43/mCherry and *ipt-*/EGFP strains, either singly, or following co-culturing for 4 h at 28 °C. Images were acquired at 100X for fluorescence channels and DIC (Nomarski). Scale bar represents 5 µm. **H**, Supplementation with exogenous cytokinin restores wildtype levels of virulence to a *ipt-* mutant of *Xoo*. TN-1 rice leaves were inoculated with BXO43, *ipt-*, or *ipt-* with injection of iP, 24 h prior to infection with *ipt-*. Lesion lengths were measured 14 days after inoculation. Error bars indicate the standard deviation of readings from 5 inoculated leaves. For all graphs, boxes capped with letters that are different from one another indicate that they are statistically different using unpaired two- sided Student’s t-test analysis (*P* ≤ 0.05). Images are representative of 3 biological replicates (**A, B, E, G**).

In order to determine if supplementation with iP would rescue the virulence deficiency of the *ipt-* strain, the strain was inoculated on rice leaves with or without injection of iP, 24h prior to infection. In the absence of iP, lesions caused by the *ipt-* strain were significantly shorter than those caused by the BXO43 strain. Supplementation with iP restored wildtype levels of virulence to the *ipt-* mutant (Fig 4H).

### The *pcrK*/*pcrR* genes are required for cytokinin sensing and virulence in *Xoo*

Recently, a cytokinin sensor named as Plant cytokinin receptor Kinase (PcrK), and its response regulator PcrR, have been identified in *Xcc* (Wang et al., 2017). Using them as a query, putative *pcrK* and *pcrR* genes were identified in the genome of *Xoo*. Microscopic analysis of the *pcrK-* and *pcrR-* strains revealed that these cells form aggregates, as compared to cells of BXO43, which remained dispersed (Fig 5A). The *pcrK-* and *pcrR-* strains also formed more biofilm as compared to BXO43 (Fig 5B-D). In order to test if exogenous cytokinin addition would rescue the aggregate formation of the *pcrK-* and *pcrR-* mutants, we added iP to actively growing cultures of the *pcrK-* and *pcrR-* mutants. However, addition of iP could neither disperse aggregate formation by cells of *pcrK-* and *pcrR-* mutants (Fig 5E) nor rescue the increased biofilm formation phenotype of these strains (Fig 5F-I). In order to check if the cytokinin secreted by BXO43 would rescue aggregate formation by *pcrK-* and *pcrR-*, we co-cultured the BXO43/mCherry strain with either the *pcrK-*/EGFP or the *pcrR-* /EGFP strains. When cultured individually, BXO43/mCherry cells were dispersed, whereas the *pcrK-*/EGFP and *pcrR-*/EGFP cells formed aggregates. On co-culturing these two mutant strains with BXO43/mCherry, we observed that the strains still showed aggregate formation (Fig 5J). This suggests that the cytokinin secreted by the BXO43 strain is unable to rescue the aggregation phenotype of the *pcrK-*/EGFP and *pcrR-*/EGFP strains.

**Figure 5.**
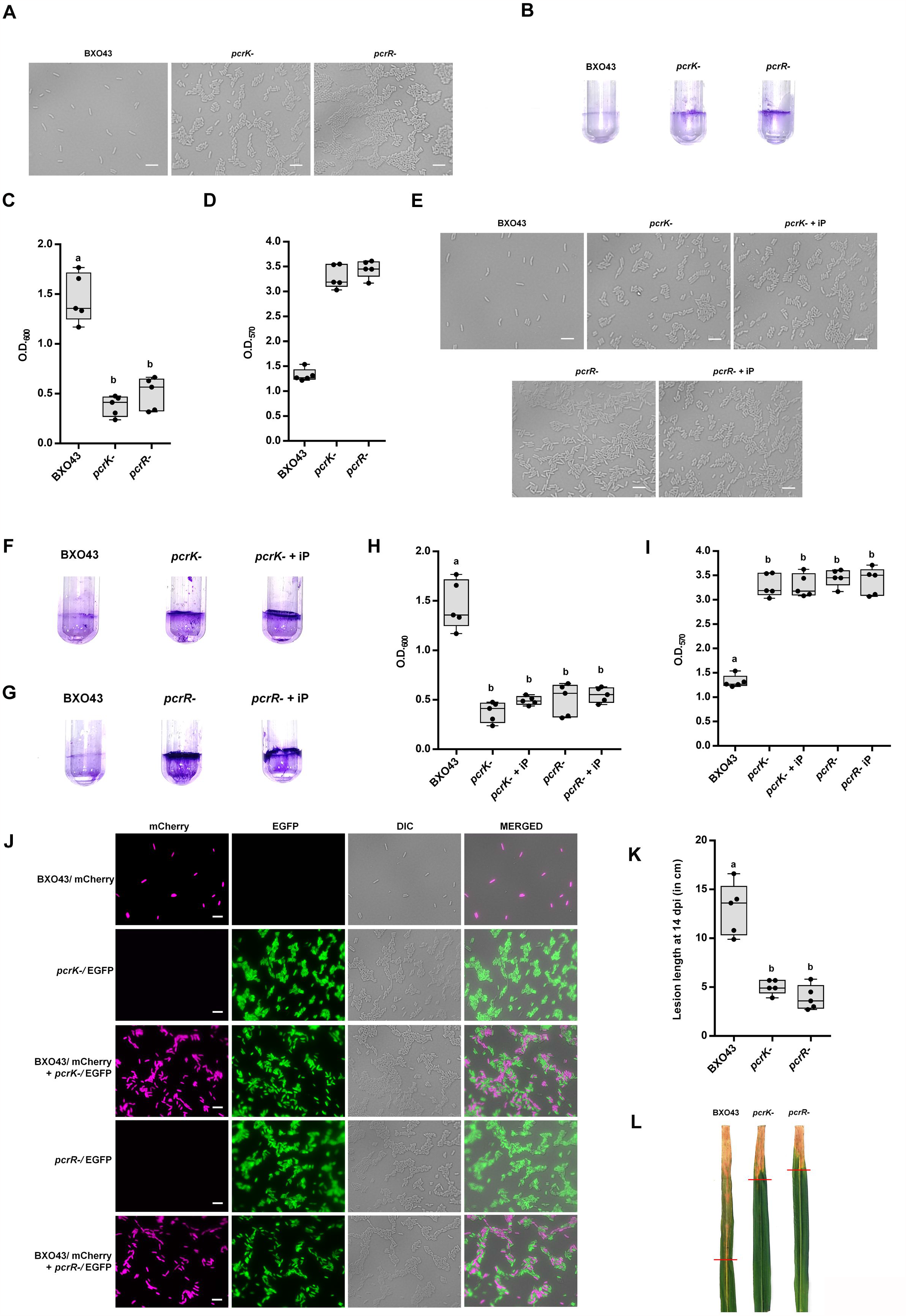
The *pcrK*/*pcrR* genes are required for cytokinin sensing and virulence in *Xoo*. **A**, Microscopy analysis was performed of the following strains: BXO43, *pcrK-* or *pcrR-*. Images were acquired at 100X in DIC (Nomarski). Scale bar represents 5µm. **B-D**. Cell attachment and static biofilm assay of *Xoo* strains BXO43, *pcrK-* or *pcrR-*. **C**, Quantification of bacterial cells in the cell suspension. Values are presented as mean absorbance (at 600 nm) ± standard deviation from 5 replicates. **D**, Quantification of bacterial cells attached to glass tubes by staining with crystal violet. Values are presented as mean absorbance (at 570 nm) ± standard deviation from 5 replicates. **E**, Microscopy analysis was performed of the following strains: BXO43, *pcrK-, pcrR-, pcrK-* + iP or *pcrR-* + iP. Images were acquired at 100X in DIC (Nomarski). Scale bar represents 5µm. **F-I**. Cell attachment and static biofilm assay of *Xoo* strains BXO43, *pcrK-, pcrR-, pcrK-* + iP or *pcrR-* + iP. **H**, Quantification of bacterial cells in the cell suspension. Values are presented as mean absorbance (at 600 nm) ± standard deviation from 5 replicates. **I**, Quantification of bacterial cells attached to glass tubes by staining with crystal violet. Values are presented as mean absorbance (at 570 nm) ± standard deviation from 5 replicates. **J**, Co- culturing of wildtype *Xoo* BXO43 with the *pcrK-* or *pcrR-* mutants does not reverse biofilm to planktonic lifestyle of *Xoo*. Microscopy analysis was performed of the BXO43/mCherry and *pcrK-*/EGFP or *pcrR-*/EGFP strains, either singly, or following co- culturing for 4 h at 28 °C. Images were acquired at 100X for fluorescence channels and DIC (Nomarski). Scale bar represents 5 µm. **K-L**, *pcrK-* and *pcrR-* mutants of BXO43 are virulence deficient. Leaves of susceptible rice TN-1 were clip inoculated with different *Xoo* strains. **K**, Lesion lengths were measured 14 days after inoculation. Error bars indicate the standard deviation of readings from 5 inoculated leaves. **L**, Virulence phenotype on rice leaves. Leaves were photographed 14 days after inoculation. For all graphs, boxes capped with letters that are different from one another indicate that they are statistically different using unpaired two- sided Student’s t-test analysis (*P* ≤ 0.05). Images are representative of 3 biological replicates (**B, F, G, J, L**).

We then proceeded to examine the virulence of the *pcrK-* and *pcrR-* strains. Leaves of 60-day old rice plants were inoculated with cultures of BXO43, *pcrK-* or *pcrR-*. As compared to BXO43, both the *pcrK-* and *pcrR-* strains showed a significant reduction in lesion length as compared to BXO43 at 14 days post inoculation, indicating that cytokinin sensing is important for complete virulence of *Xoo* (Fig 5K, L).

### XopQ regulates the biofilm to planktonic lifestyle switch in *Xoo*

In order to gain a greater understanding of the regulatory role of XopQ, RNA-sequencing (RNA-seq) analysis was performed to assess differential gene expression between BXO43 and the *xopQ-* strain. In the absence of XopQ, there were a total of 757 differential expressed genes (DEGs), of which 328 were down-regulated and 459 were up-regulated. Differential expression of 10 such genes was validated by RT-qPCR (Appendix Table S4, Appendix Figure S5A, b). GO analysis revealed the abundance of bacterial motility, bacterial flagellar assembly and protein transport related functional categories (Appendix Figure S5C). Further, KEGG pathway analysis using the genome annotation of *Xoo* strain BXO1 (Midha, Bansal et al., 2017), revealed “biofilm formation”, “bacterial chemotaxis” and “flagellar assembly” related pathways to be upregulated in the *xopQ-* mutant. Upregulation of the pathway for biofilm formation was consistent with our observation that the *xopQ-* mutant forms more biofilm. In this regard, upregulation of *cheA, cheB, cheV, cheW, cheY* and *mcp* genes was noteworthy in the *xopQ-* mutant. Silencing of *mcp, cheB*, and *cheV* by RNAi has earlier been shown to lead to deficiencies in adhesion, chemotaxis, flagellar assembly and motility (Huang, Wang et al., 2017). This may lead to increased cellular adhesion, similar to what is observed in the *xopQ-* mutant. Furthermore, the KEGG analysis revealed components of type IV secretion system to be up-regulated in the *xopQ-* mutant and previous results indicate this secretion system is involved in promoting biofilm formation (Cenens, Andrade et al., 2020, Elhenawy, Hordienko et al., 2021, Seifert, 2017). Notably, components of the type III secretion apparatus were amongst down-regulated pathways in the *xopQ-* mutant (Appendix Table S5). We observed the reduced expression of protein components of the type III secretion system (T3SS) and multiple type III effectors in the *xopQ-* mutant, which could be rescued by the addition of iP (Appendix Figure S5D). This suggests that expression of these proteins is regulated by cytokinin and may explain the reduced virulence of the *xopQ-* mutant. Expression of the T3SS has previously been shown to be repressed in biofilm-growing bacteria (Kuchma, Connolly et al., 2005).

## Discussion

The production of the phytohormone cytokinin by various phytopathogenic bacteria to modulate host physiology and virulence is well known. However, there are no reported examples of bacterial cytokinin production modulating bacterial physiology or social behaviour. Here we report the production of cytokinin by the rice pathogen *Xoo* and present evidence that cytokinin production controls the switch between biofilm and planktonic states. An important initial step in colonization during bacterial infection is adhesion. In *Xanthomonas*, this has been shown to involve the expression of multiple virulence factors, which includes surface appendages such as flagellum and type IV pili, which are required for the colonization of host tissues (Huang et al., 2017, Qi, Huang et al., 2020). However, post-colonisation spread in the host plant requires a switch from biofilm to planktonic lifestyle for an effective spread *in-planta* (Appendix Figure S6). We propose that bacterial cytokinin regulates this switch in *Xoo* and promotes virulence.

Our results indicate that the *Xoo* type III effector XopQ is a phosphoribose-hydrolase and can convert iPMP to iP. XopQ and its orthologs have earlier been shown to be required for full virulence, and immune response modulation (Deb, Ghosh et al., 2020, Deb, Gupta et al., 2019, Giska, Lichocka et al., 2013, Gupta et al., 2015, Li, Chiang et al., 2013, Li, Yadeta et al., 2013, Teper, Salomon et al., 2014) (Appendix Figure S7). *Arabidopsis* transgenic plants expressing the *Pseudomonas* ortholog HopQ1 show suppression of Flg22-induced defense responses by attenuating flagellin receptor FLS2 expression in a cytokinin dependent manner (Hann et al., 2014). HopQ1 has been predicted to catalyse the last step in the production of cytokinin, converting iPMP to the active cytokinin iP. In flowering plants, this step is catalysed by the cytokinin riboside 5′-monophosphate phosphoribohydrolase (LONELY GUY/ LOG) class of enzymes, which catalyze the formation of active cytokinin species from cytokinin ribosides (Kurakawa, Ueda et al., 2007). Recently, the only homolog of this enzyme from the unicellular green microalga *Chlorella* was shown to be a cytokinin-activating enzyme (Nayar, 2021).

The *Xoo* genome encodes a homolog of the IPT protein that is predicted to catalyze the committed step in cytokinin production. The *ipt* mutant is defective in cytokinin production as well as virulence and exhibits the same aggregation phenotype as the *xopQ-* mutant. These phenotypes are reversed by cytokinin supplementation. The estimation of cytokinin levels in *Xoo* revealed that the levels of iP was almost 100-fold more as compared to trans-zeatin (tZ). This is similar to what is observed in the cyanobacterium *Nostoc* (Frébortová, Greplová et al., 2015). *Nostoc* was also shown to have a complete cytokinin synthesis machinery, with a conserved isopentenyl transferase (IPT) protein and a cytokinin dehydrogenase (CKX) protein (Frébortová et al., 2015). Our transcriptome data indicates upregulation of the biofilm formation pathway as well as type IV bacterial secretion pathway in the *xopQ-* mutant as compared to BXO43. This might explain why the *xopQ-* mutant has an enhanced biofilm formation phenotype. A reduced expression of the type III secretion system, and multiple type III effectors is observed in the *xopQ-* mutant, suggesting that this could also lead to the reduced virulence of the *xopQ-* mutant. How might cytokinin be sensed by *Xoo*? Our studies suggest that mutations in either the predicted cytokinin receptor kinase (PcrK) or the response regulator PcrR of *Xoo* results in phenotypes that are akin to those of the *xopQ-* and *ipt-* mutants, suggesting that these proteins could be involved in cytokinin sensing by *Xoo* (Fig 5).

Interestingly, the *ipt* gene and the 4777 bp region encompassing it are present in some but not all members of the genus. The presence of this gene cluster in some but not all *Xanthomonas* species, the atypical codon usage pattern and the presence of a tRNA gene near the *ipt* cluster are consistent with the possibility that this gene cluster may have been inherited by horizontal gene transfer. A large number of bacteria have genes that are predicted to encode phosphoribose-hydrolase and isopentenyl transferase activities and a few have been shown to produce cytokinin (Samanovic et al., 2015, Seo & Kim, 2017). Thus, cytokinins are produced by a number of different bacteria. Our results demonstrate for the first time that endogenously produced cytokinin regulates physiological activities in bacteria. We postulate that cytokinin may be an important signalling molecule in a number of bacterial species. Furthermore, we suggest that the origins of cytokinin as a signalling molecule may be rooted in bacteria and that this role may have been subsequently elaborated upon in the plant kingdom.

## Materials and methods

### Bacterial strains, plasmids, media and growth conditions

The plasmids and bacterial strains used in this study are listed in Appendix Table S1 and Appendix Table S2 respectively. The *Xoo* strains were grown at 28 °C in peptone-sucrose (PS) media (Daniels et al., 1984). *Escherichia coli* strains were grown in Luria–Bertani (LB) medium at 37 °C. Antibiotics were added at the following concentrations in the media: rifampicin: 50 µg/mL; kanamycin: 50 µg/mL (*E. coli*), 15 µg/mL (*Xoo*); spectinomycin: 50 µg/mL; gentamicin: 10 µg/mL.

### Microscopy

To monitor cell morphology, overnight grown *Xoo* strains were harvested, concentrated and immobilized on a thin agarose pad of 2 % agarose and visualized under a Zeiss AxioImager microscope in DIC (Nomarski optics) mode. For co-culturing of strains, the overnight grown cultures (BXO43/mCherry and *xopQ-*/EGFP or *ipt-*/EGFP) were adjusted to equal cell density, mixed and incubated at 28 °C for 4 h. For assays involving the addition of cytokinin, the overnight grown cultures were adjusted to equal cell density, and 20 nM iP was added to the secondary culture, which was grown at 28 °C for 24 h.

### Confocal microscopy for biofilm visualisation

To monitor biofilm formed by the *Xoo* strains, BXO43, *xopQ-* and *ipt-* strains were transformed with the EGFP-expressing plasmid pMP2464 (Appendix Table S1) (Stuurman, Pacios Bras et al., 2000), grown overnight at 28 °C and normalized to an O.D._600_ of 0.1. 15 ml of the culture was taken in a 50 ml tube and a sterilised glass slide was introduced into it. This was kept stationary at 28 °C for 72 h. To evaluate biofilm formation, the slide was washed gently with MQ water to remove loosely adhering cells, and further fluorescent Z-stacked images were acquired of 0.38 µm to measure overall attachment and biofilm levels at the air-culture interface on the slide at 63X in a LSM880 confocal microscope. The Zen software was used to plot the GFP signal intensity profile for the Z-stacked images for biofilm thickness and Imaris software was used to process the images for surface 3D visualisation.

### Quantitative cell attachment and static biofilm assays

*In-vitro* biofilm formation was visualised and quantified. The strains were grown overnight at 28 °C and normalized to an O.D._600_ of 0.1 in PS media in glass tubes. This was incubated for 4 days at 28 °C without shaking. In order to quantify cells remaining in planktonic state, the culture was decanted carefully, and OD_600_ reading was taken. The glass tube was washed three times with MQ gently to remove any non-adhering cells. The resultant biofilm was further stained with a 0.1% crystal violet solution, at room-temperature for 30 min. Following this, the stain was removed, excess stain washed off, and the tubes were imaged. For quantification of biofilm, the crystal violet stain was solubilized using a combination of 40 % methanol and 10 % glacial acetic acid. Data was collected in the form of the O.D._570_ of the elution.

### *In-vitro* enzyme assay

Phosphoribose-hydrolase activity for XopQ and its mutants was performed with purified proteins by using the substrate N6-(2-isopentenyl) adenine-9-riboside-5′-monophosphate (OlChemIm Ltd., Olomouc, Czech Republic; Cat. No: 001 5043) as described previously (Hann et al., 2014) with modifications. The assay mixture contained purified 1pM enzyme supplemented with various concentrations of the substrate, in a 200µl reaction buffer (50mM HEPES, 100mM NaCl, 10mM imidazole pH 7.0 containing 1mM dithiothreitol and 5mM CaCl_2_). The reaction was performed for 5 s at 37 °C, and terminated by the addition of NaOH to a final concentration of 0.1 N. Product formation was measured as change in absorption at 280 nm. 6-(γ,γ-dimethylallylamino) purine (Sigma; Cat. No: D5912-5G) was used for calculation of standard curve.

### Virulence assays

60-day-old rice plants of the susceptible rice ‘Taichung Native’ (TN-1) were used for assays for virulence. *Xoo* strains were grown to saturation and inoculated by dipping scissors into bacterial cultures of O.D._600_=1 and clipping the tips of rice leaves. Lesion lengths were measured at 14 days after inoculation and expressed as the mean lesion length with standard deviation.

In order to study the effect of exogenous supplementation of cytokinin, 10nM isopentenyl adenine or trans-zeatin was injected into the midvein of 60-day old TN-1 rice leaves. 24 h post injection, pin-prick inoculation of the respective *Xoo* strains was done 1 cm above the point of injection. Lesion lengths were measured at 14 days after inoculation and expressed as the mean lesion length with standard deviation.

### Estimation of cytokinin

Bacterial strains were grown to saturation, cell pellet and supernatant were separated and lyophilised, and cytokinin was extracted by methanol/formic acid/water (15/0.1/4 v/v/v), using the internal standards *trans*-[^2^H_5_]zeatin and [^2^H_6_]isopentenyl adenine (OlChemIm Ltd., Olomouc, Czech Republic). The extracts were purified using an C_18_ RP SPE column and analysed using a 6500+ Qtrap system coupled with ultra-performance liquid chromatography using a Zorbax C18 column.

### Codon Usage Pattern

Codon Usage Pattern (CUP) was calculated for each gene to estimate the frequency of codon usage for different amino acids as described previously (Patil & Sonti, 2004), using “The Sequence Manipulation Suite” webtool (Stothard, 2000). Briefly, eight amino acids (Glycine, Valine, Threonine, Leucine, Arginine, Serine, Proline and Alanine) were selected, which have at least four synonymous, and the percentage of codons that end with G or C was calculated for each amino acid and gene. The first group was chosen to include housekeeping genes that encode proteins which participate in various essential functions in *Xoo*. These genes encode: BXO1_013815 (TonB-dependent siderophore receptor), BXO1_013910 (*Xanthomonas* adhesin like protein), BXO1_006505 (*rpfF*), BXO1_016165 (shikimate dehydrogenase) and BXO1_019245 (secreted xylanase). The LPS cluster, which was earlier shown to have come in *Xoo* by horizontal gene transfer (Patil & Sonti, 2004), was taken as a control group. This group consisted of five genes of the LPS cluster: BXO1_014260 (*smtA*), BXO1_014255 (*wxoA*), BXO1_014250 (*wxoB*), BXO1_014240 (*wxoC*) and BXO1_014235 (*wxoD*).

### Western blotting

Bacterial cultures were grown to saturation, pelleted and analysed for the presence of XopQ protein. Cells were lysed by sonication and total protein supernatants were isolated after centrifugation at 14,000 rpm for 15 min at 4 °C to remove cellular debris. Equal amounts of isolated protein supernatants were further used for Western blotting. The XopQ protein was detected using anti-XopQ antibodies raised in rabbit. Immunoblotting was carried out using ALP conjugated to anti-rabbit immunoglobulin G secondary antibody (Sigma Aldrich; A3687). Equal loading of protein in the different samples was shown using Coomassie blue staining of gels.

### Global transcriptome analysis using RNA-seq

Total RNA was sequenced at the NGC facility of CDFD, Hyderabad, with RNA isolated from the cell pellets of *Xoo* strains (BXO43 and *xopQ-*) grown to an O.D._600_=1 in PS media. Quality of the RNA was checked on Agilent TapeStation 4200. Ribosomal RNA (rRNA) depletion was carried out using the NEBNext® rRNA Depletion Kit (Bacteria), and library preparation was carried out using NEBNext® Ultra™ II Directional RNA Library Prep Kit for Illumina®. Prepared libraries were sequenced on Illumina Nextseq2000 (P2 200 cycle sequencing kit) to generate 60M, 2×100bp reads/sample. The sequenced data was processed to generate FASTQ files. Differential gene expression analysis was conducted on the generated data using STAR-featureCounts-DEseq2 pipeline. A false discovery rate (FDR) ≤ 0.05, and |log_2_ of the fold changes| ≥1 was considered for differentially expressed genes. Gene Ontology enrichment analyses were performed with PANTHER using the Gene Ontology Resource (2021, Ashburner, Ball et al., 2000, Mi, Muruganujan et al., 2019) and the pathway analyses were performed using KEGG database.

### RNA isolation and gene expression analysis

For RNA isolation, bacterial cultures were grown to O.D._600_ =1, pelleted and RNA was extracted using Macherey-Nagel RNA isolation kit according to the manufacturer’s instructions, which included on-column digestion of genomic DNA. 5 μg of total RNA was reverse transcribed into cDNA using EcoDryTM Premix (Clontech, Mountain View, CA, USA) according to the manufacturer’s instructions using random hexamer primers. Synthesized cDNA was diluted 5-fold and then used for semi-quantitative RT-PCR with 35 cycles of amplification. 16S rRNA was used as an internal control. The cDNA was analysed for the presence of *xopQ* (600 bp N-terminal fragment) and *ipt* (750 bp full-length gene) transcripts. Absence of genomic DNA was confirmed using a set of primers from a non-coding unique region of the genomic DNA.

Transcript analysis of genes found to be differentially expressed by RNA-seq in the *xopQ-* mutant as compared to BXO43, was carried out by reverse transcriptase-quantitative polymerase chain reaction (RT-qPCR). RT-qPCR of selected genes (Appendix Table S4) was performed using gene-specific primers using Power SYBR Green PCR Master Mix (Thermo Fisher Scientific) in BioRad CFX384 Real-Time PCR System (Hercules, California, United States). Relative expression was calculated with respect to BXO43. The fold change was calculated using 2-ΔΔCt method (Livak & Schmittgen, 2001). Expression of 16S rRNA gene was used as internal control.

### Bioinformatic analysis of IPT protein

Multiple sequence alignment of the IPT protein from various bacterial strains was carried out using T-Coffee multiple sequence alignment server (Expresso) (Notredame, Higgins et al., 2000). The GenBank ID of the IPT homologue in *Xanthomonas oryzae* pv. *oryzicola* BLS256 is AEQ96873, in *Xanthomonas oryzae* pv. *oryzae* PXO99A is ACD58327, in *Xanthomonas albilineans* is WP_012917043, in *Xanthomonas translucens* is WP_053834798, in *Xanthomonas theicola* is WP_128421291, in *Agrobacterium rhizogenes* is WP_080705458, in *Ralstonia solanacearum* is WP_119447925, in *Ensifer psoraleae* is WP_173514402, in *Agrobacterium vitis* is WP_070167542, in *Pseudomonas savastanoi* is AGC31315, in *Pseudomonas amygdali* is WP_081007393, in *Rhizobium tumorigenes* is WP_111221635, in *Sinorhizobium* sp. PC2 is WP_046120136, in *Agrobacterium tumefaciens* is QTG17184 and in *Pseudomonas psychrotolerans* is WP_193755078.

For phylogenetic analysis of the IPT protein, a phylogenetic tree was constructed based on the sequence of IPT proteins from bacteria, plants, fungi, and Archaea using the MEGA X software (Kumar, Stecher et al., 2018). Briefly, iterative searching for IPT protein was performed using position-specific iterated BLAST (PSI-BLAST) method in NCBI (National Centre for Biotechnology Information) (Altschul, Madden et al., 1997). Phylogenetic tree analyses was conducted in MEGA X software using Maximum Likelihood method based on Le Gascuel 2008 model (Le & Gascuel, 2008). The tree with the highest log likelihood (−56444.14) is shown.

## Author Contributions

SD, HKP and RVS conceived and designed the experiments. SD performed all the biological assays, and wrote the manuscript. CK and RK performed the cytokinin estimation. AK performed the genomic bioinformatic analysis. PG assisted in generation of the *pcrK-* and *pcrR-* mutants. SD, HKP, SC, GJ, PP and RVS analyzed the data, and finalized the manuscript, which was approved by all the authors. HKP, GJ and RVS contributed reagents/materials.

## Availability of data and material

The GEO code for the RNA-sequencing data generated for this study is GSE179029.

